# Spectral flow cytometry reveals metabolic heterogeneity in tissue macrophages

**DOI:** 10.1101/2022.05.26.493548

**Authors:** Graham A Heieis, Thiago A Patente, Tamar Tak, Luís Almeida, Bart Everts

## Abstract

Tissue-macrophage populations are constituted by a mosaic of phenotypes, yet new methods are needed to link metabolic status to the range of phenotypes *in vivo*. We therefore designed a high-dimensional panel for spectral flow cytometry to investigate the heterogeneity of tissue macrophage metabolism at steady-state, and their metabolic adaptation in response to infection. Distinct metabolic profiles were observed between tissue macrophages from different peripheral organs, as well as within populations from a specific site. As such, our data show multiple metabolic states in macrophages corresponding to relative stages of maturity in both the peritoneal cavity and small intestine. Immune perturbation with helminth infection further showed that peritoneal macrophages acquire an overall highly metabolically active profile, whereas responding intestinal macrophages displayed minimal changes in their metabolic phenotype. Thus, we demonstrate that high-dimensional, flow-based analysis is an exciting method to interrogate the metabolic heterogeneity and dynamics of tissue-macrophage populations.

## Introduction

Macrophages (Mphs) support immune defense, tissue integrity and homeostatic processes in multiple biological systems. Embryonically seeded mphs persist as long-lived inhabitants within tissues, maintaining numbers through local proliferation (Nobs and Kopf, 2021). Conversely, adult monocytes make varying contributions to the mph population at steady state depending on the tissue (Ginhoux and Guilliams, 2016). Inflammation, however, can expedite the replacement of resident mphs due to cell death and parallel recruitment of monocytes. Provided the diverse tissue environments they inhabit, mphs require distinct signals for differentiation, and are phenotypically discernable, across various distal sites (Nobs and Kopf, 2021).

Accumulating evidence argues strongly that metabolic reprogramming is critical for mph function and residence (Wculek et al., 2021), yet few reports have dissected the active metabolic phenotypes of *in vivo* mphs outside of transcriptomic data. However, evidence up to this point indicates different metabolic niches permit different responses to immune and pathogenic stimuli (Scott et al., 2022; Svedberg et al., 2019). The caveat to *in vivo* metabolic characterization of mphs is the difficulty in isolating sufficient numbers of mphs for metabolic studies, and maintaining confidence that cellular metabolism does not lose fidelity as a result of removing them from their tissue-niche.

Metabolic flow or mass cytometry targeting nutrient transporters and enzymes has recently been pioneered as a highly promising alternative to analyze immune metabolism at the single-cell level (Ahl et al., 2020; Geeraerts et al., 2021; Hartmann et al., 2021). These methods for metabolic analysis have yet to define the tissue-specific metabolic states, or responses, of mphs. We therefore adapted a similar approach here to investigate metabolic features of mphs *ex vivo* using spectral flow cytometry, a relatively new technology endowed with several advantages, including high-throughput acquisition and the ability to resolve lowly-expressed antigens. Our data reveal dynamic metabolic states between early monocyte-derived versus resident mphs, marked diversity of tissue mph metabolism within and across peripheral organs, and site-specific changes in metabolism during helminth infection. These findings emphasize the advantage of spectral flow-based analysis for metabolic targets, and serve as a foundation to further explore unique metabolic properties and heterogeneity in tissue-resident macrophage populations in health and disease.

## Results

### *in vitro* differentiated macrophages acquire divergent metabolic phenotypes

We made a selection of nutrient transporters and metabolic enzymes of core metabolic pathways based on previous studies, and of known relevance to mph function (Ahl *et al*., 2020; Covarrubias et al., 2020; Hartmann *et al*., 2021) (Figure 1A). Analysis of murine bone marrow-derived mphs (BMDMs) treated with inflammatory stimuli (LPS+IFNγ), to induce a classically activated ‘M1’ phenotype, demonstrated increased features of glycolysis and the pentose-phosphate pathway, as well as an altered tricarboxylic acid (TCA) cycle. This expression profile was consistent with metabolic flux data, which showed an increased extracellular acidification rate (ECAR), representative of glycolysis (Figure S1A-C). In contrast, IL-4-stimulated BMDMs with an alternatively-activated ‘M2’ phenotype, maintained high expression of lipid and amino acid transporters relative to LPS+IFNγ stimulated mphs, in line with a high oxygen consumption rate (OCR) detected by Seahorse (Figure S1A-C). Exploiting the cross-reactive nature of the antibodies targeting metabolic markers, we assessed the metabolic state of GM-CSF or M-CSF differentiated human monocyte-derived mphs (hMDMs), which promote pro- and anti-inflammatory features in hMDM, respectively (Fleetwood et al., 2007). Several parallels were observed between BMDMs and hMDMs, including higher GLUT1, PKM and SDHA expression in inflammatory GM-CSF compared to regulatory M-CSF hMDM (Figure S1D-E). These data show that analysis of metabolic targets detects expected differences in both human and murine mph metabolism *in vitro*.

**Figure 1.**
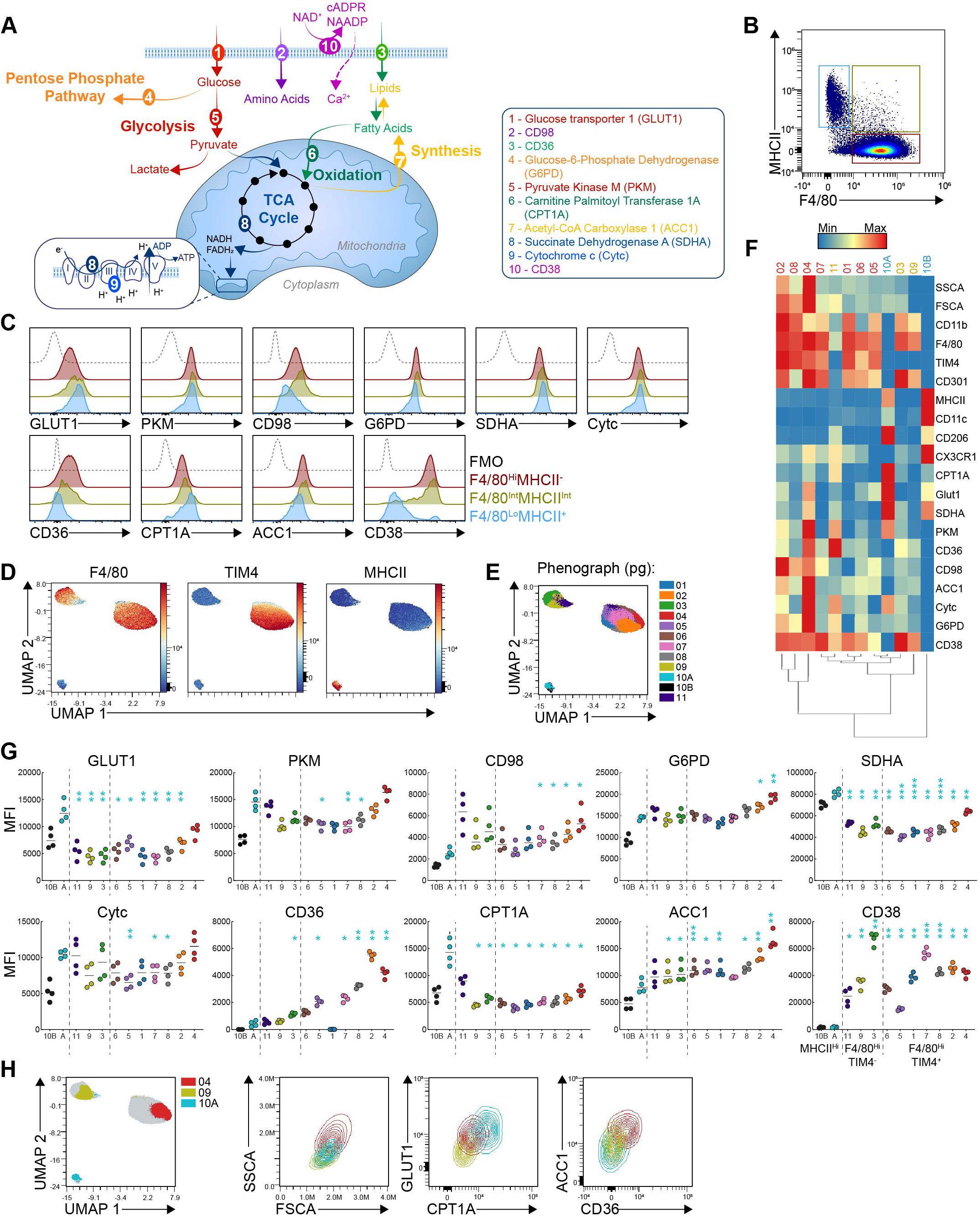
Macrophages display metabolic heterogeneity within the peritoneal cavity. (**A**) Schematic of metabolic targets for flow-based analysis. (**B**) Live, single CD45^+^CD11b^+^ cells (CD3^-^B220^-^ Ly6C^-^Ly6G^-^Siglec F^-^) from the peritoneal cavity. (**C**) Histograms for metabolic targets in peritoneal Mph populations shown in (B). (**D**) UMAP dimensional reduction, (**E**) phenograph clusters and (**F**) heatmap of Lin^-^CD11b^+^ cells using phenotypic (F4/80, CD11b, CD11c, MHCII, TIM4, CX3CR1, CD301, CD206) and metabolic markers. (**G**) MFI of metabolic targets in clusters from (F). (**H**) Contour plots of clusters pg04, 05 and 10A showing differential cell size and metabolic expression. Analysis performed on cells pooled from 4 mice from one representative out of 3 experiments. p<0.05(*), 0.01(**), 0.001(***), 0.0001(****), from RM one-way ANOVA. Asterisks match the color of the cluster to which the comparison is made..

### Dynamic metabolic states can be identified in peritoneal cavity macrophages

An additional advantage of spectral flow cytometry is the ability to define autofluorescence (AF) as a new channel, allowing superior resolution of fluorophores. However, the emission of varying AF spectra can become an obstacle for spectral “unmixing” due to heterogenous populations or high AF intensity, as exhibited by digested tissues and mphs. We developed a workflow to define AF spectra for optimal unmixing of all immune cell populations, which we successfully applied in a 27-channel panel (excluding AF channels)(Figure S2). We first characterized total peritoneal exudate cells (PEC), which resulted in similar metabolic expression patterns between myeloid and lymphoid populations as has been reported for human blood leukocytes (Ahl *et al*., 2020; Hartmann *et al*., 2021) (Figure S3A-C).

We then initially specifically investigated the metabolic heterogeneity of peritoneal mphs. Population analysis of recruited immature (MHCII^Hi^F480^Lo^), intermediate (MHCII^Int^F4/80^Int^) and mature resident (MHCII^-^F4/80^Hi^) mphs revealed divergent metabolic profiles (Figure 1B-C). In particular, fatty acid-transporter CD36, amino acid-transporter CD98, and the ecto-NADase CD38 were up-regulated in resident cells. By contrast, the glucose transporter GLUT1 and the mitochondrial fatty acid transporter CPT1A were highest in MHCII^Hi^F480^Lo^ mphs. To interrogate these differences further, dimensional reduction and clustering were applied to total Lin^-^CD11b^+^ cells using a combination of phenotypic and metabolic markers. Uniform Manifold Approximation and Projection (UMAP) revealed three distinct mph populations, defined mainly by MHCII, F4/80 and TIM4 (Figure 1D). Phenograph clustering identified several F4/80^Hi^ mph clusters (pg01-09), but only a single MHCII^Hi^ cluster (pg10), which appeared to consist of at least two populations based on size (FSC-A) and MCHII expression (Figure 1E, S3D). Thus cluster pg10 was split into two clusters defined as pg10A (MHCII^+^FSC-A^Hi^) and pg10B (MHCII^++^FSC-A^Lo^). The expression pattern of pg10A aligned with a well-described phenotype for immature peritoneal mphs, whereas pg10B contains both early mph precursors as well as CD11b^+^ DCs (Bain et al., 2016).

Metabolic expression across mph clusters revealed a pronounced elevation in CPT1A and GLUT1, as well as the dual Kreb’s cycle/electron transport chain enzyme SDHA, in cluster pg10A that was notably lower in F480^Hi^TIM4^-^ clusters (Figure 1F-G). In contrast, metabolic marker expression was overall similar between TIM4^-^ (pg03,09,11) and TIM4^+^ (pg01, 02, 05-8) mphs, of which the latter appeared to separate largely by variable expression of TIM4, CD301 and CD38 (Figure 1F-G and S3E). Interestingly, within the TIM4^+^ population, there was heterogeneity in the expression of ACC1,which facilitates the primary step in fatty acid synthesis. The clusters with highest ACC1 expression (pg02 and pg04) also exhibited greater cell size and granularity (FSC-A & SSC-A), a trait of resident mphs (Louwe et al., 2021), compared to TIM4^-^ and MHCII^Hi^ mphs (Figure 1F-H, S3E). Altogether, these data illustrate phenotypic and metabolic characterization by spectral flow cytometry can reveal heterogeneity within mph populations. Furthermore, dynamic metabolic protein expression suggests monocyte-derived differentiation is associated with altered glucose and lipid metabolism in the peritoneal cavity.

### Resident macrophages of different tissues display heterogenous metabolic phenotypes

To extend our characterization across mph populations from different tissues, we assessed metabolic expression in resident mphs from the PEC, lung, liver and lamina propria of the colon and small intestine (CLP and SILP, respectively) (Figure 2A-D, S4). Liver Kupffer cells (KCs) exhibited higher expression of nearly all targets, indicating they are more metabolically active in comparison to counterparts from other tissues (Figure 2A-C). Resident large peritoneal mphs (LPMs, F4/80^Hi^TIM4^+^) had similarly high expression of most markers relative to KCs, although KCs remained unique in their high expression of CD36, CD98 and CD38. In contrast, mphs from either the small or large intestine, as well lung alveolar mphs (AlvMs), displayed relatively metabolically quiescent profiles. In AlvMs, specifically CD36, CD98 and G6PD expression were elevated over intestinal mphs, while SDHA and Cytc expression was comparatively low. Clustering according to only metabolic expression yielded discrete populations for each tissue, with the exception of the small intestine and colon, which overlapped (Figure 2B-C). Of note, peritoneal mphs incubated at 37^°^C alongside digesting tissues maintained comparable expression from matching samples stored immediately on ice (Figure S4D), indicating tissue processing steps did not significantly impact analyzed metabolic parameters.

**Figure 2.**
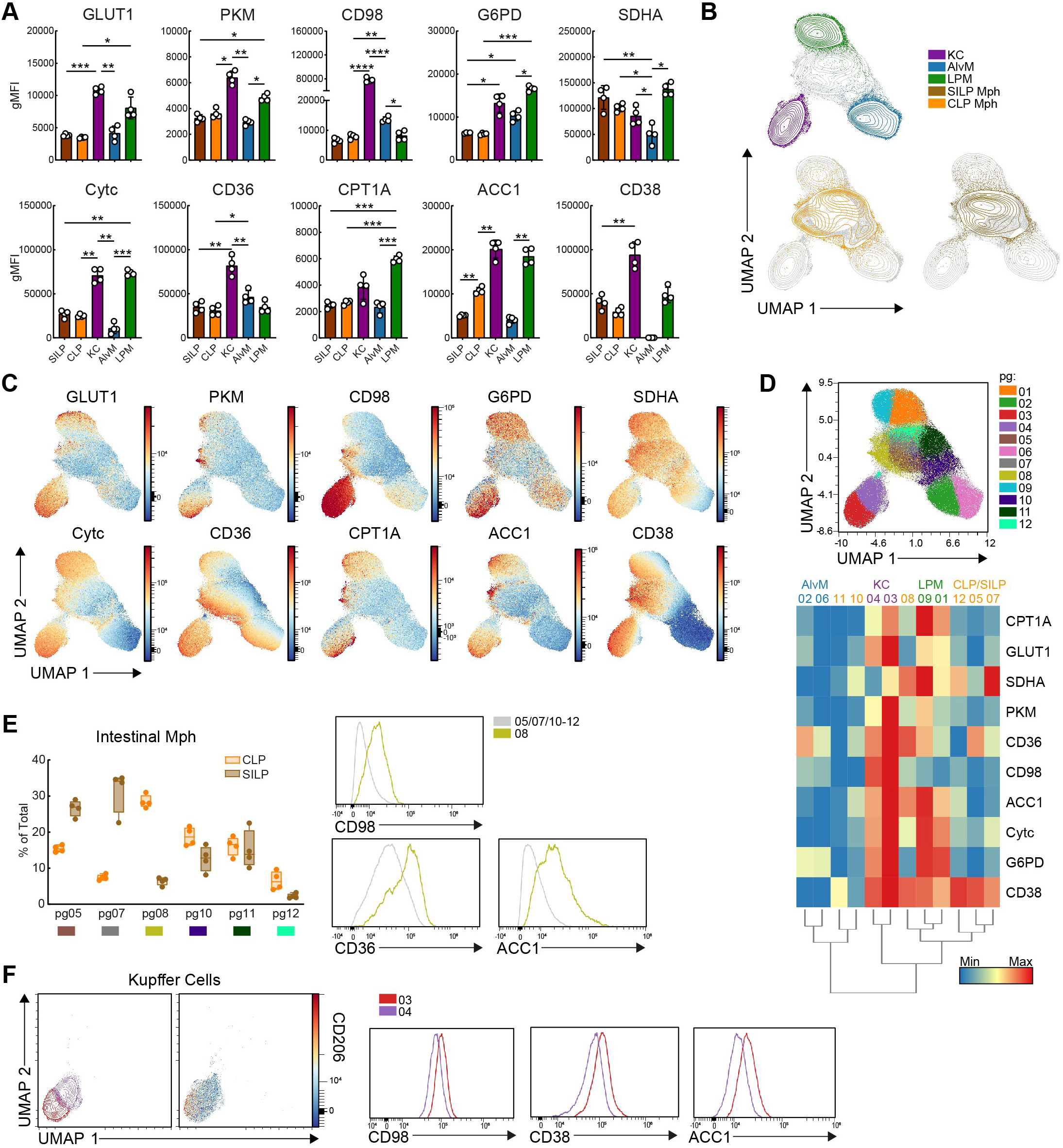
Tissue-tropic metabolic phenotypes across resident macrophage populations. Intestinal (CD11b^+^CD64^+^MHCII^Hi^Ly6C^-^), Liver (CD11b^Lo/Mid^CD64^+^TIM4+), alveolar (CD11c^+^SigF^+^) and peritoneal (F480^Hi^TIM4^+^) mph were gated in FlowJo and exported for analysis in OMIQ. (**A**) gMFI of metabolic targets across tissue-mph populations. (**B**) UMAP of tissue mph generated according to metabolic target expression, and (**C**) the relative distribution of metabolic targets. (**D**) Phenograph clustering and resulting heatmap using only metabolic targets of tissue mph from (A). (**E**) Relative abundance of clusters for CLP and SILP mphs and histograms displaying differential expression between the most abundant cluster in the colon and all other clusters. (**F**) Phenograph clustering of liver mphs distinguishing clusters resembling KC1 and KC2 using only metabolic protein expression and corresponding histograms. Analysis done with pooled data from 4 mice, representative of 2 experiments (n=2-4). p<0.05(*), 0.01(**), 0.001(***), 0.0001(****), ns=not significant, calculated by RM one-way ANOVA.

Multiple clusters were identified in mphs from each tissue with intestinal mphs showing the greatest diversity (Figure 2D, E). Despite the overlap of colonic and small intestinal mphs, there was appreciable variability in the abundance of each cluster between sites (Figure 2E). For instance, cluster pg08 contained the majority of colonic mphs, and displayed higher expression of CD98, CD36 and ACC1 in relation to other intestinal clusters (Figure 2E).

KCs have recently been discovered to contain two populations, termed KC1s and KC2s, of which the latter is defined by CD206 expression (Bleriot et al., 2021; De Simone et al., 2021). Clustering our data yielded two KC populations (pg03, pg04) that displayed differing levels of CD206, in accordance with one cluster resembling KC2s (pg03) (Figure 2F). The putative KC2 cluster also had increased expression of ACC1, CD98 and CD38 over KC1s (Figure 2F), supporting and extending transcriptomic data that KC2s exhibit particularly elevated fatty acid synthesis, amino acid catabolism and adenosine metabolism (Bleriot *et al*., 2021). Together, these data reveal tissue-specific, and tissue intrinsic, heterogenic metabolic patterns in mph populations.

### Intestinal infection drives tissue-dependent metabolic responses in macrophages

Macrophage metabolism is commonly assessed using polarizing stimulations *in vitro*, yet how metabolic responses to these stimuli translate to tissue-mphs remains largely unanswered. Alternative mph activation in response to IL-4, for instance, causes metabolic reprogramming towards glycolysis (Huang et al, 2016), lipolysis (Huang et al., 2014) and glutaminolysis (Liu et al., 2017), but this has yet to be physiologically validated *in vivo. Heligmosomoides polygyrus* (Hp) is an enteric parasite that leads to alternative activation in the small intestinal lamina propria (SILP) as well as the peritoneal cavity, despite strict localization to the proximal small intestine (Filbey et al., 2014). Hence, we characterized metabolic and polarization profiles of mphs from the PEC and SILP (duodenum) of naïve and infected mice (Figure 3A, S4C).

**Figure 3.**
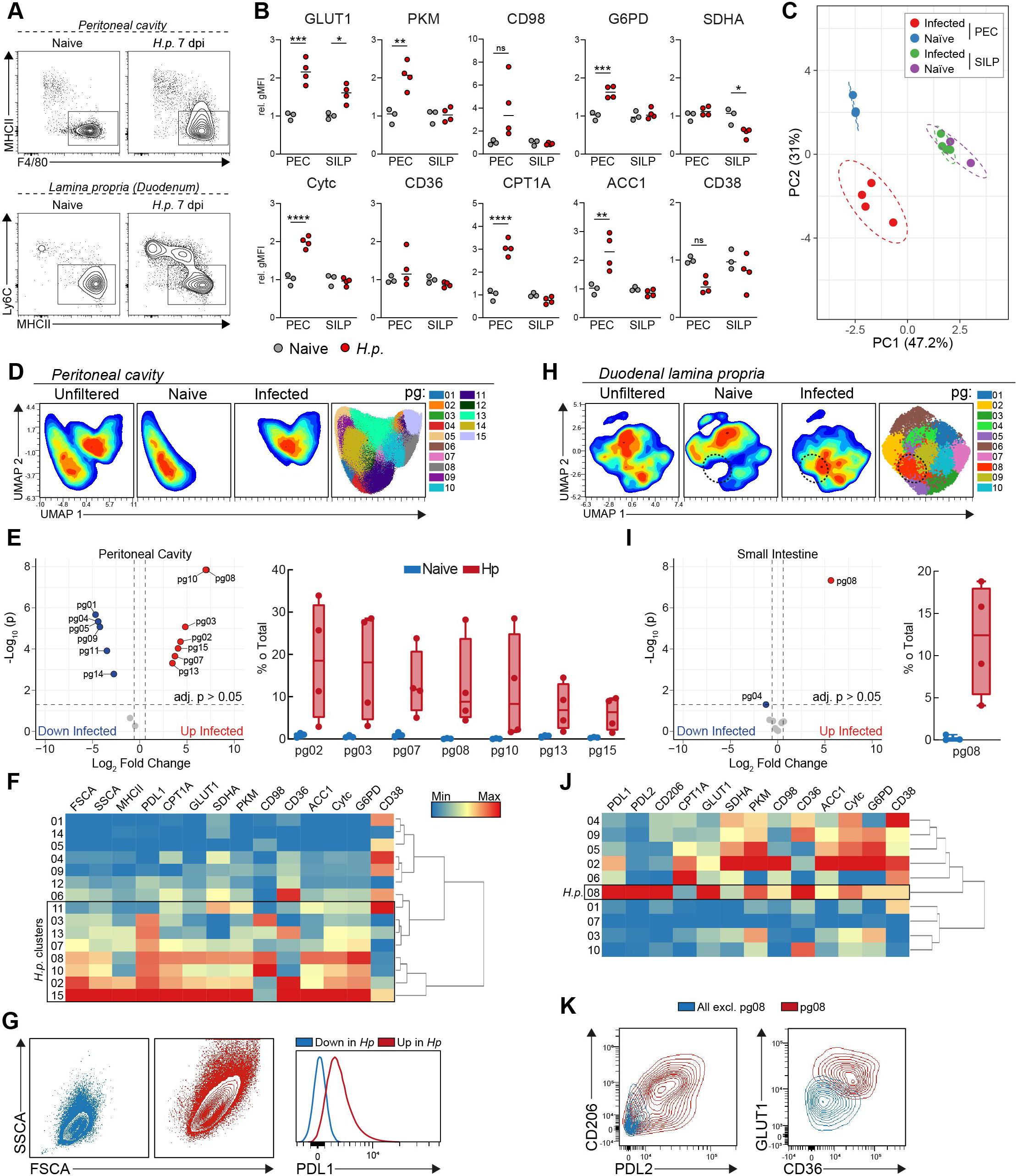
Macrophages respond metabolically to intestinal helminth infection *in vivo*. Mice were left naïve or infected with *H. polygyrus* L3 larvae and sacrificed 7 days after infection. (**A**) Representative staining for peritoneal Mph (Lin^-^F480^+^CD11b^+^) and lamina propria monocytes/mph (Lin^-^CD64^+^CD11b^+^) from the proximal 15cm of the small intestine. (**B**) Relative gMFI of metabolic targets in mature Mph (PEC: F4/80^Hi^MHCII^Lo^, SILP: MHCII^Hi^Ly6C^-^), normalized to the average gMFI of Mph from uninfected mice. (**C**) Principle-component analysis using gMFI of mph metabolic protein expression. (**D**) UMAP and phenograph clustering of peritoneal Mph. (**E**) EdgeR analysis revealing clusters of differential abundance between naïve and infected mice. (**F**) Heatmap showing expression across peritoneal Mph clusters. (**G**) Forward and side scatter, and PD-L1 expression of indicated peritoneal Mph clusters. (**H-J**) Analysis of SILP Mph as described for PEC mphs in D-F. (**K**) Expression of alternative activation markers, CD36 and GLUT1 from SILP cluster pg08. Infection vs naïve comparison are representative of one experiment with 3 and 4 naïve and infected mice, respectively. Data from naïve group are representative of 2 experiments. p<0.05(*), 0.01(**), 0.001(***), 0.0001(****), ns=not significant, calculated by two-way ANOVA.

Infection led to global increases of metabolic enzyme expression in PEC mphs, while only changes in GLUT1 and SDHA were observed in mphs from the infected SILP, which were increased and decreased, respectively (Figure 3B). Principal component analysis of the expression of metabolic targets further showed that peritoneal mphs diverge according to infection status, whereas mphs from the SILP consistently grouped together, despite observed changes in GLUT1 and SDHA (Figure 3B,C).

The stark differences in metabolic responses between tissues prompted deeper characterization of mphs from either site. Clustering of peritoneal mphs based on metabolic target expression similarly showed strong divergence in response to Hp, with significantly different abundance in 13/15 clusters between groups (Figure 3D-E). Clusters prevalent during Hp infection correlated with both a metabolically and an immunologically active phenotype, with an overall increased expression of metabolic markers, MHCII and PDL1, as well as cell size (Figure 3F-G). Interestingly, CD38 has been recently identified as a marker of inflammatory mphs *in vitro* (Covarrubias *et al*., 2020), which was found to be decreased in peritoneal mphs during Hp infection (Figure 3F), and thus may be negatively correlated with alternative activation *in vivo*.

Corresponding analysis of SILP mphs again revealed substantial overlap between naïve and infected groups (Figure 3H). However, one cluster (pg08) appeared during Hp infection that was altogether absent in the naïve intestine (Figure 3I). This cluster expressed CD206, PDL1 and PDL2, indicating alternative activation (Figure 3J,K). When comparing pg08 to all other SILP clusters, the most dominant metabolic changes observed were increased GLUT1 and CD36 expression (Figure 3J,K). Notably, increases of metabolic targets observed in PEC Mphs were absent in intestinal mphs during infection, including CPT1A. Therefore, simultaneous analysis of mphs from the PEC and SILP during Hp infection reveals metabolic responses differ according to tissue localization.

### Metabolic adaptation during macrophage differentiation in the naïve and infected duodenum

Mph differentiation from adult monocytes in the intestine is associated with progressive changes in Ly6C, MHCII, CX_3_CR1, and CD11c (Bain et al., 2013; Desalegn and Pabst, 2019), whereas TIM4^+^ mphs have been identified as embryonically seeded (Shaw et al., 2018). We found that helminth infection promoted the former process, also known as the monocyte waterfall, as visualized by differential Ly6C and MHCII expression (Bain et al., 2014)(Figure 3A). We therefore questioned the metabolic dynamics of macrophage differentiation in the small intestine. As shown earlier (Figure 3B), only GLUT1 and SDHA expression differed between mphs from naïve and Hp infected mice using bulk population analysis (Figure 4B). CD36, CD38 and CD98 expression was associated with phenotypically mature mphs in the SILP, while GLUT1 trended towards higher expression in intermediate mphs (similar to the naïve PEC, Figure 2G). Interestingly, and in contrast to peritoneal mphs which displayed higher CPT1A expression in immature mphs (Figure 2G), CPT1A was highest in mature mphs in the intestine of naïve mice (Figure 4B).

**Figure 4.**
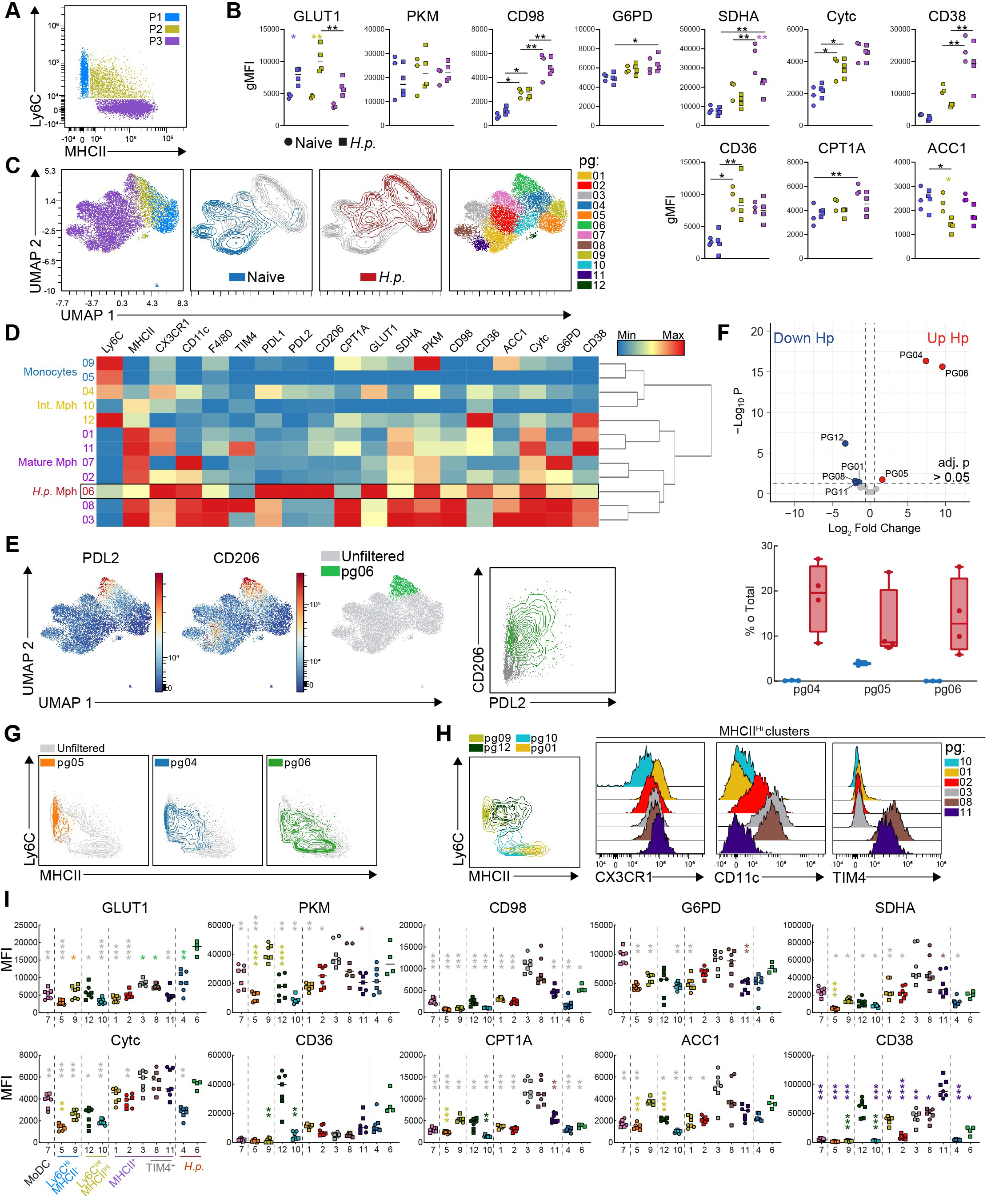
Metabolic transitions during the monocyte waterfall in the naïve and *H. polygryus* infected duodenum. (**A**) Total CD11b^+^CD64^+^ leukocytes from the proximal small intestine, pooled from naïve and Hp infected mice. (**B**) Metabolic marker gMFI of manually gated Ly6C^Hi^MHCII^-^, Ly6C^Int^MHCII^Int^ and Ly6C^-^MHCII^Hi^ populations from naïve and Hp infected mice. (**C**) UMAP and phenograph clustering of CD11b^+^CD64^+^ cells using metabolic and lineage targets. (**D**) Heatmap of clusters from (C). (**E**) Identification of CD206 and PD-L2 expressing Mph and corresponding phenograph cluster. (**F**) EdgeR analysis showing clusters statistically more prevalent during Hp infection, and (**G**) their Ly6C and MHCII expression. (**H**) Identification of relative cluster maturity, or origin, using Ly6C, MHCII, CX3CR1, CD11c and TIM4 expression. (**I**) MFI of metabolic targets in clusters identified in (C) arranged according to expression patterns from (H). Analyzed data are from the same experiment as Figure 2. p<0.05(*), 0.01(**), 0.001(***), 0.0001(****), using (B) two-way or (I) RM one-way ANOVA. Asterisks match the color of the cluster to which the comparison is made..

In accordance with the expansion of Ly6C^Hi^ and intermediate monocytes, unbiased analysis of CD11b^+^CD64^+^ immune cells using metabolic and lineage markers led to separation between naïve and infected samples (Figure 4C). A cluster of alternatively activated cells (pg06), expressing more GLUT1 and CD36, was similarly identified as from earlier analysis (Figure 3J,K & 4D,E). These mphs resembled recently differentiated monocytes, with residual Ly6C and comparatively low MHCII expression (Figure 4D,F,G). Indeed, all clusters more abundant during infection displayed traits of differentiating monocyte-derived mphs at various stages (Figure 4F, G).

Next, we predicted the relative maturity of intestinal mphs along a developmental timeline according to the aforementioned markers (Figure 4H). Early differentiating CX_3_CR1^+^CD11c^Lo^ mphs (pg01, 02) maintained a relatively low metabolic profile, comparable to transitioning monocytes. However, one cluster of monocytes (pg09) displayed particularly high PKM and ACC1 expression, whereas a fraction of intermediate monocytes (pg12) uniquely expressed high levels of CD36 and CD38. Amongst Ly6C^-^MHCII^Hi^ mphs, increasing CD11c expression paralleled increases in most metabolic targets, except CD36 and CD38. These data suggest maturation of intestinal mphs is associated with an increased metabolic phenotype, but may involve transitioning through a series of dynamic metabolic states from monocytes. Interestingly, we identified two populations of TIM4^+^ cells. The first (pg11) showed marked similarity to CX_3_CR1^Hi^CD11c^Lo^ monocyte-derived mphs (pg01) with the exception of higher CD38 expression. The second TIM4^+^ cluster (pg08) displayed a phenotype similar to that of cluster pg03, characterized by high CD11c and metabolic marker expression (Figure 4H, I). TIM4^+^ cells are reported to have minimal turnover from monocytes (Shaw *et al*., 2018); thus, our data may indicate that metabolic profiles of different mph populations in the small intestine are shaped by inflammatory cues, rather than ontogenetic differences.

## Discussion

Despite the widely recognized limitations surrounding *in vitro* experiments, a prolonged lag in pushing towards *in vivo* studies has hindered advances in mph immunometabolism (Van den Bossche and Saraber, 2018). Flow-based analysis of metabolic enzyme expression has been demonstrated as a technique to investigate metabolic features of immune cell populations from blood (Ahl *et al*., 2020; Hartmann *et al*., 2021) or tumors (Geeraerts *et al*., 2021; Levine et al., 2021). Here, to the first of our knowledge, we present the use of spectral flow cytometry for high dimensional analysis of tissue mphs, adapting ‘met-flow’ to interrogate mph metabolism. Our investigation focused on murine mphs, however the cross-reactivity of the antibodies, as demonstrated with human MDMs, means our panel is readily usable for interrogating the metabolic profiles of mphs from human tissues.

Each tissue-niche has its own controlled metabolic microenvironment (Wculek *et al*., 2021). Mphs are therefore predicted to have complementary metabolic wiring to support their habitation in any particular tissue (Bleriot *et al*., 2021; Bonnardel et al., 2019; Davies et al., 2017; Svedberg *et al*., 2019). AlvMs exist in a highly lipid-concentrated space and depend on PPARγ, a key regulator of lipid uptake and oxidation, for development (Schneider et al., 2014). Accordingly, next to KCs, AlvMs exhibited the greatest CD36 expression, and had the lowest ACC1 expression of all mph populations. In accordance with KC dependance on LXRs (Bonnardel *et al*., 2019), which can promote fatty acid synthesis and glycolysis via SREBP activation (Assmann et al., 2017; Bonnardel *et al*., 2019; Sakai et al., 2019; Schultz et al., 2000), we observed comparatively high ACC1, GLUT1 and PKM in KCs. CD98 was also highly expressed in KCs compared to other mphs. Little is known of amino acid metabolism in KCs, however the recently identified KC2 subset had enriched transcripts for amino acid pathways over KC1s, implying some functional significance in liver homeostasis (Bleriot *et al*., 2021). Our data also align with recently published work demonstrating small intestine mphs are metabolically less active than those from the large intestine, specifically in regards to amino acid and lipid metabolism(Scott *et al*., 2022).

Here we found notable differences between early monocyte-derived mphs and phenotypically mature mphs. Most strikingly in the peritoneal cavity, immature MHCII^Hi^ mphs displayed highly elevated expression of enzymes involved in glycolysis and fatty-acid oxidation, that was not evident in F480^Hi^ cells. Similar, but less evident, trends were also observed in the small intestine, in which intermediate monocytes displayed transiently increased GLUT1, PKM and CPT1A, but not to the level observed in CD11c^Hi^MHCII^Hi^ mphs of the same tissue. In line with these findings, increased PPARγ has been linked to monocyte-derived peritoneal mph differentiation (Gautier et al., 2012). It was more recently demonstrated that CCR2-controlled deletion of CPT1A abrogates differentiation of monocyte-derived mphs following cardiac transplant (Zhu et al., 2022). Together with our data, these studies indicate a rapid shift, and requirement, for glycolysis and fatty acid oxidation in the process of mph development from monocytes. Furthermore, the lipid receptor/transporter ApoE is a key feature of TIM4^+^ mphs (Bain et al., 2020), and here we show increased CD36 and ACC1 expression, strongly arguing for a metabolic switch towards lipid accumulation in resident mphs. As dead cell clearance via efferocytosis is a core function of peritoneal mphs, it is tempting to speculate that lipid accumulation facilitates membrane organization and organelle synthesis to support this process (Lee et al., 2018).

We further show here that metabolic responses of mphs to helminth infection varied dramatically between sites. Intestinal mphs in particular showed little metabolic responsiveness. A population resembling alternatively-activated mphs was identified based on altered GLUT1 and CD36 expression, however, our findings that these mphs expressed low levels of Ly6C and MHCII suggest these cells were newly recruited entrants into the tissue, rather than a metabolic adaptation of resident mphs. Functional and metabolic hypo-responsiveness to type 2 stimuli has been reported in lung AlvMs, which was attributed to limited glucose availability in alveolar spaces (Saumon et al., 1996; Svedberg *et al*., 2019). It is improbable that glucose concentrations would be limiting in the proximal small intestine (Ferraris et al., 1990), thus host or parasite-derived immuno-suppressive factors may instead dampen metabolic responses in the intestine (Ip et al., 2017; Schridde et al., 2017). This notion would be consistent with observations that intestinal mphs display a general resistance to stimulation (Bain *et al*., 2013). The Hp-driven increases in GLUT1 and CD36 in intestinal mphs support previously ascribed needs for glucose and fatty acids of mphs in the context of type 2 immunity (Huang *et al*., 2014; Huang et al., 2016). Conversely, fatty acid oxidation is frequently cited as a requirement for alternative activation (Wculek *et al*., 2021), yet we see no evidence of altered CPT1A expression in activated, duodenal mphs during Hp infection. The role of fatty acid oxidation for *in vitro* polarization has already been questioned (Divakaruni et al., 2018; Nomura et al., 2016), and here we make further observations in support of this argument *in vivo*.

In conclusion, despite the well-appreciated importance and heterogeneity of tissue mphs, few studies have attempted to characterize their metabolic diversity, instead relying on *in vitro* systems, RNA-sequencing or inferences from transgenic models. With our work combining spectral flow cytometry and metabolic protein analysis, we have generated a foundation and valuable tool for continued exploration of the *in vivo* aspects of mph metabolism.

## Acknowledgements

This work was supported by the LUMC, by the Leiden University Fund/Schild-de Groen Fonds, (www.luf.nl) awarded to G.A.H, and by an NWO Vidi grant (#91719349) awarded to B.E. We would like to acknowledge LUMC Flow Core Facility operators, in particular IJsbrand Reyneveld, for the continual maintenance and trouble-shooting of the Cytek Auroras. Many thanks to our colleagues in LUMC Parasitology for their continual scientific discussions and critical reading of the work.

## Author Contributions

G.A.H and B.E. conceived the study. G.A.H designed, performed and analyzed experiments with the assistance of T.A.P., T.T. and L.A. The manuscript was written by G.A.H and B.E.

## Declaration of Interests

No competing interests to declare by the authors

## Main Figure Legends

## Methods

### Mice and infections

C57BL/6 mice were used, bred in-house and maintained under SPF conditions, or purchased from Envigo. Mice were used between 8-16 weeks of age. Experiments were complete in compliance with the Guide for the Care and Use of Laboratory Animal Research, and with approval from the Dutch Central Authority for Scientific Procedures on Animals (CCD; license number: AVD1160020198846). *H. polygyrus* infection was given by oral gavage of 200 L3 larvae. Mice were sacrificed on day 7 post-infection. Hp larvae were generously provided by Prof. Rick Maizels (University of Glasgow, UK).

### Murine Bone Marrow Derived Macrophages

Femurs and tibias of mice were collected in RPMI, surface sterilized with ethanol and flushed with cold HBSS using a 25g needle and syringe. Clumps and fragments were removed by aspirating up and down with the syringe followed by filtering through a 40μM strainer into a 50ml tube. Bone marrow cells were centrifuged at 300g for 5min, resuspended and counted in cold RPMI. Cells were adjusted to concentration in RPMI containing 10% heat-inactivated FCS, 100U/ml penicillin, 100μg/ml streptomycin, 50μM 2-mercaptoethanol and 10% supernatant of M-CSF producing L929 cells. 5ml containing 2×10^6^ cells were plated in non-culture treated 6-well plates. Cultures were supplemented on day 3 or 4 with 5ml of complete RPMI containing 20% L929 supernatant. Cells were harvested using accutase on day 7, replated in 24-well plates with 3-5×10^5^ cells/well and stimulated in 1ml for 18-24 hours with LPS (100ng/ml) and IFNy (50ng/ml) or IL-4 (20ng/ml).

### Human Monocyte Derived Macrophages

Monocytes were isolated from buffy coat and differentiated into MDM. Blood was diluted 1:3 in Hanks’ Balanced Salt Solution (HBSS) and placed on 12 mL of Ficoll to obtain mononuclear cells. The solution was centrifuged at 400 x g for 30 minutes at RT, without brake. The resulting mononuclear cell layer was removed, placed in a new tube containing HBSS, and centrifuged at 300g for 20 min. The cells were washed twice more and monocytes were isolated using CD14 MACS beads (Miltenyi) according to the manufacturer’s recommendations, routinely resulting in a monocyte purity of >95%. To differentiate into macrophages 2×10^6^/well of monocytes were cultured in 6 well plates using RPMI supplemented with 10% FCS, 100 U/mL of penicillin, 100 μg/mL streptomycin, 2 mM of glutamine, and either 20 ng/mL rGM-CSF (Invitrogen) or 20ng/ml rM-CSF for 6 days. Cultures were supplemented on day 2/3 with an equal volume of 2x concentrated cytokines.

### Seahorse Metabolic Analysis

1×10^5^ murine BMDM were plated for stimulation in an XFe96 well Seahorse plate and stimulated overnight. Media was replaced, after 2x washes with PBS, with 80μl XF assay made from base RPMI supplemented with 10mM glucose, 2mM glutamax, and 2mM sodium-pyruvate, and incubated in a non-CO_2_ 37 °C incubator for 1 hour. Immediately before analysis, an additional 95μl of XF media was added to all cells. As cells were incubating injected compounds were diluted in XF media and added to the hydrated cartridge, after which the cartridge was immediately loaded into the Seahorse for calibration. Oligomycin = 2μM, FCCP = 1.5μM, Rotenone/Antimycin = 0.75μM each (1.5μM total).

### Tissue immune cell isolation

Peritoneal exudate was obtained by injection of 5ml cold PBS containing 2% FCS and 2mM EDTA into the exposed abdomen of sacrificed mice, which was subsequently withdrawn after ∼20 seconds of gentle agitation, and kept on ice until counting and plating.

Published protocols were adapted for isolation of leukocytes from the small-intestine (Webster et al., 2020) and colon (including caecum) (Platt et al., 2010). Both tissues were placed on PBS-soaked paper towel, opened longitudinally (after removal of Peyer’s patches from small intestine), and intestinal contents were scraped off using a metal spatula. Opened intestines were washed vigorously by shaking in a 50ml tube containing 15ml of Ca/Mg-free HBSS containing 2mM EDTA (wash media), then cut into 1-2cm pieces and stored in 10ml of wash media on ice until further processing. To remove mucous and strip epithelial cells, tissues were shaken vigorously in 50ml tubes with 10ml pre-warmed wash media, strained through 250μM Nitex, placed back in tubes containing 10ml fresh wash media, and incubated for 20 min at 37°C, shaking at 200rpm. Small intestines were washed a total of 3 times, colons a total of 2 times. Small intestine digest: 10ml RPMI containing 10% FCS, 1mg/ml collagenase VIII, 40U/ml DNAse I for 15-20min (or until tissue is not quite completely digested). Colon digest: 10ml RPMI containing 10%FCS, 1mg/ml collagenase IV, 0.5mg/ml collagenase D, 1mg/ml dispase II, 40U/ml DNAse I for 25-30min. Digestion was stopped by immediate addition of cold RPMI containing 10% FCS. Digested intestines were filtered through a 100μm strainer, spun at 400g for 5min, resuspended in FACS buffer (2% FCS, 2mM EDTA) and filtered a second time through a 40μM strainer, spun, resuspended and stored on ice for counting and plating.

Liver digestion was also adapted from previously published protocols (Lynch et al., 2018). Livers were taken and placed in cold RPMI, minced with scissors and immediately transferred to tubes containing 10ml digest buffer (same as colon). Minced livers were digested for 25-30min at 200rpm, 37deg, with additional manual shaking vigorously by hand every 5-8 min. Digested livers were filtered through 100um strainers, and topped-up with cold RPMI containing 10% FCS up to 50ml, spun at 300g for 5min. Supernatant was removed, and cells were washed a second time with 30ml FACS buffer. The remaining pellet was treated with red-blood cell lysis buffer, washed and counted for staining.

lungs were placed in a 2ml tube with cold PBS, removed and minced in a new 2ml tube, before direct addition of 1.5ml digest buffer consisting of 1mg/ml collagenase IV and 40U/ml DNAse I. Lungs were shaken for 30min before storing on ice, mashing through 100μM filter and washing with 10ml cold RPMI with 10% FCS. Lung homogenate was red-blood cell lysed, washed, and passed through a 40μm strainer before counting and plating.

### Staining procedure for spectral flow cytometry

A complete list of antibodies and dilutions is shown in Table S1. Intracellularly stained metabolic targets were conjugated in-house using the corresponding kit according to the manufacturer’s protocol. For *in vitro* experiments, all cells from a single well were stained after transferring to 96-well V-bottom plate. For tissues, 1-2×10^6^ cells (numbers kept consistent within experiments) were stained in V-bottom plates in 50μl for each step. Cells were pre-stained with viability dye and Fc-block in PBS for 15min on ice. If required, subsequent live surface-staining was performed for certain targets in FACS buffer for 30 min on ice (Table S1) before fixation. Cells were fixed with eBioscience Foxp3 fixation/permeabilization staining kit according the recommended protocol, before staining intracellular targets, and intracellular Fc-block, in 1x permeabilization buffer for 1 hour at 4 deg. After intracellular staining, cells were washed 2x with permeabilization buffer. Remaining surface targets were stained in FACS buffer for 30min at 4°C All staining was done in the appropriate buffer containing 10x Brilliant Stain Buffer Plus and 20x TrueStain Monocyte Block. Single-color and unstained references were prepared in parallel for each tissue, with controls being stained simultaneous to their corresponding step for the fully stained samples. Cells were acquired on a Cytek Aurora 5-laser spectral flow cytometer. Acquired samples were unmixed using SpectroFlo version 3 and analyzed with FlowJo and/or OMIQ.

### Spectral Unmixing

Where possible, single-stained references were generated using cells from matching tissues, otherwise beads were used. References were stained in parallel to fully stained samples to follow the exact same protocol, including fixation and washing steps. References were cleaned in SpectroFlo to eliminated AF and increase clarity of the positive peaks. The “Unstained” was exported as AF negative cells. AF was defined using the CD45 reference control, gating CD45(R8)^+^ cells, and assessing AF in the UV, violet and/or blue channels, gating and exporting AF populations before re-importing as a new channels. The “Autofluorescence as a fluorescent tag” box was additionally selected for unmixing. In some cases minor compensation adjustments were made using the SpectroFlo software.

### OMIQ analysis and workflow

Populations were first gated in FlowJo and exported (Figure 2-4, total Mph; Figure 5, total live CD45+) for uploading into OMIQ. Parameters were scaled using conversion factors ranging from 6000-20000, gated according to Mph population(s) of interested, and subsampled using a maximum equal distribution across groups/tissues. After sub-sampling, UMAP was performed using the indicated parameters, followed by phenograph clustering (k=100). Data was further analyzed with EdgeR to determine significantly different clusters, or by generating heatmaps with Euclidean clustering. OMIQ was also used to generate scatter, contour and histogram plots. Graphs were generated using GraphPad Prism, showing geometric fluorescence intensity data exported from FlowJo or median fluorescent intensity as determined by OMIQ.

### Statistical analysis

Ex vivo data were tested for normality using the Shapiro-Wilk test. Differences between MFI of OMIQ-defined clusters were analyzed by a repeated measures one-way ANOVA and Tukey test correction for multiple comparisons. In the case of multiple group comparisons, a two-way ANOVA was used. Where indicated data were normalized by the mean value of the naïve group. For in vitro experiments, stimulated cells were normalized to untreated (BMDM) or M-CSF treated cells from the same donor. For BMDM a RM one-way ANOVA was employed, with Geisser-Greenhouse correction and Dunnett Test to correct for multiple comparisons. Human MDM were analyzed using Wilcoxon matched-pairs signed rank test. *p* values < 0.05 were considered significant (*p < 0.05, **p < 0.01, ***p < 0.001, ****p < 0.0001) and statistical analyzes were performed using GraphPad Prism v.9.0.

**Table S1:**
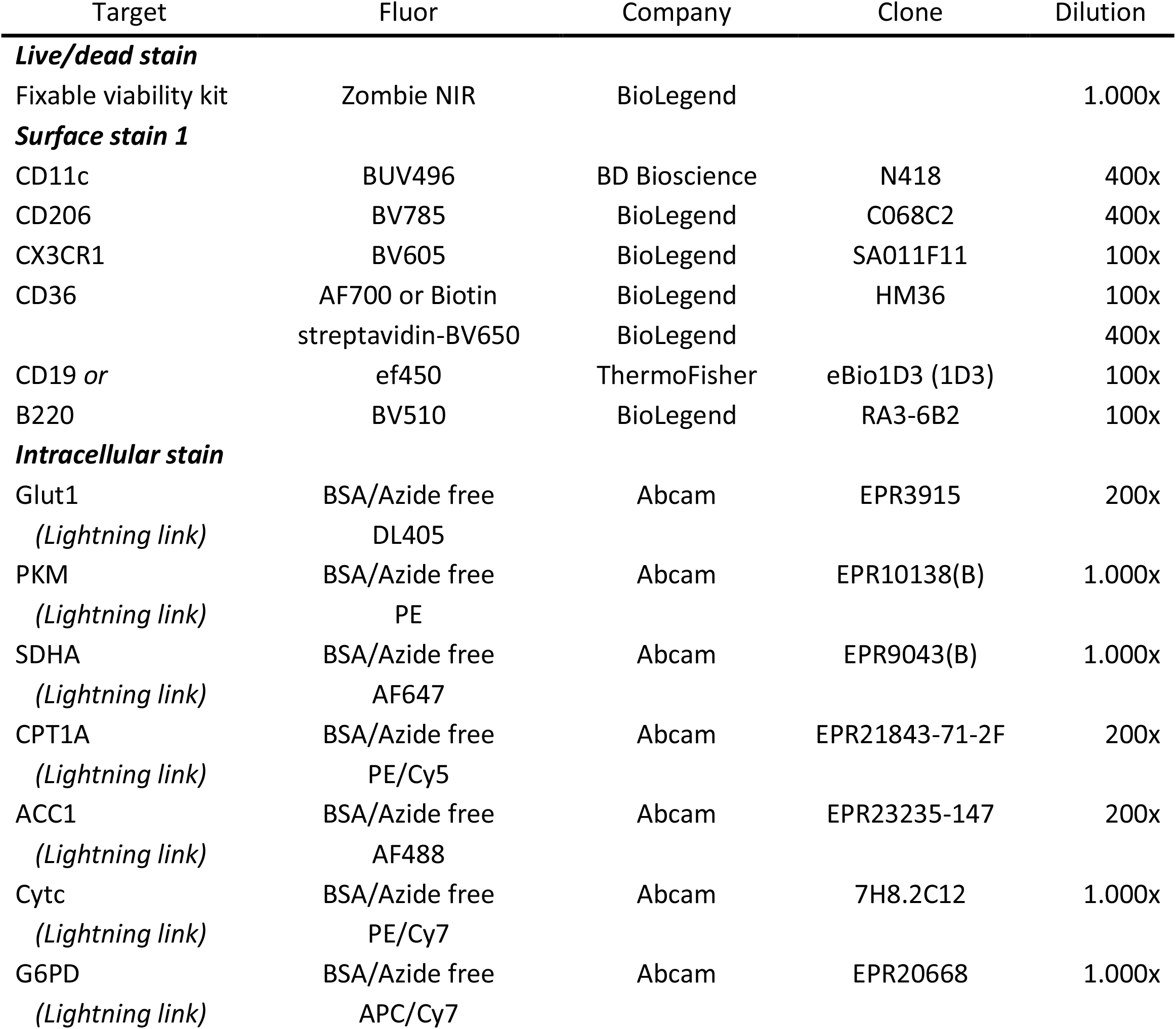
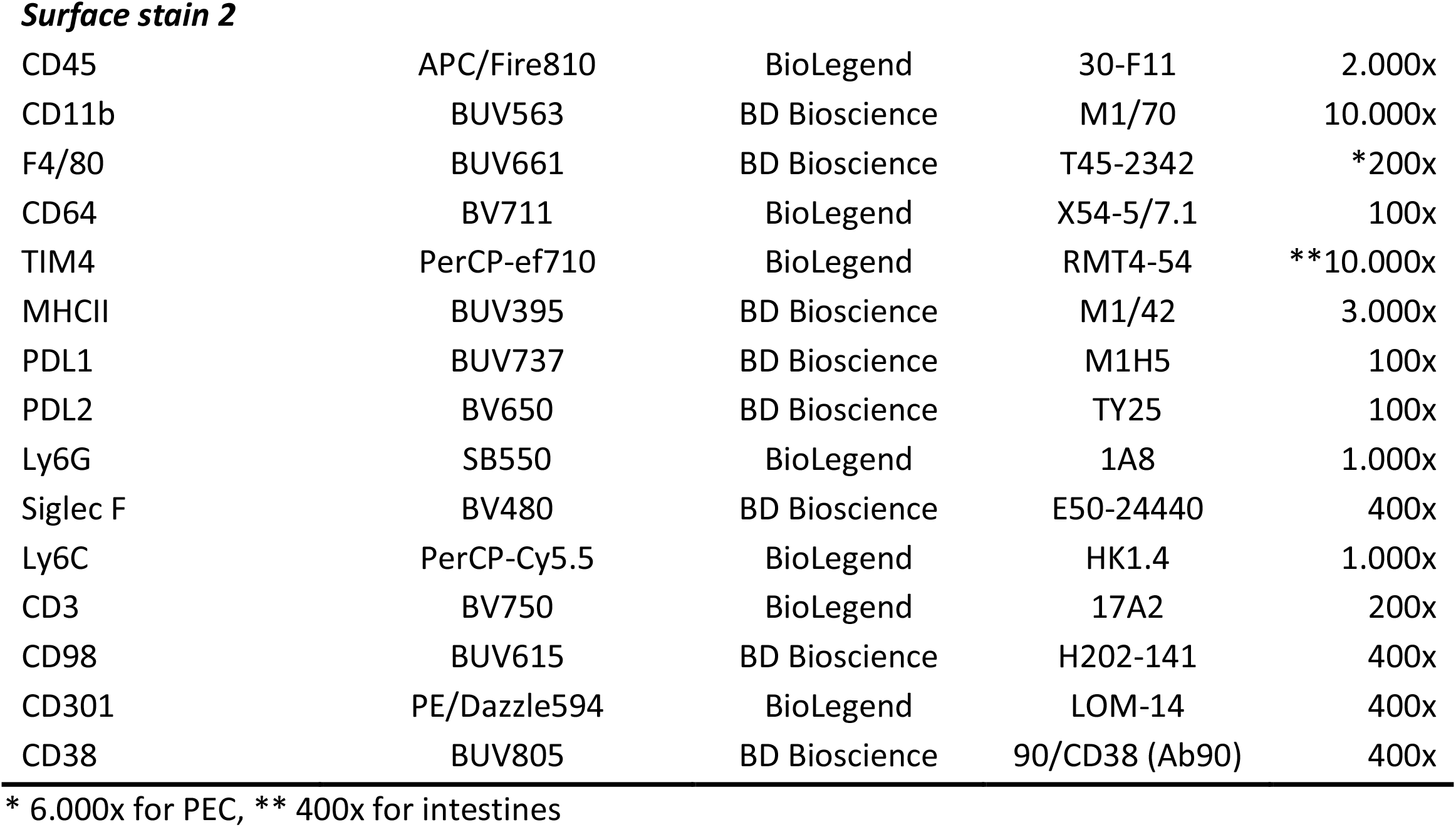
Flow cytometry reagents and staining order.

| Target | Fluor | Company | Clone | Dilution |
| --- | --- | --- | --- | --- |
| ***Live/dead stain*** |  |  |  |  |
| Fixable viability kit | Zombie NIR | BioLegend |  | 1.000x |
| ***Surface stain 1*** |  |  |  |  |
| CD11c | BUV496 | BD Bioscience | N418 | 400x |
| CD206 | BV785 | BioLegend | C068C2 | 400x |
| CX3CR1 | BV605 | BioLegend | SA011F11 | 100x |
| CD36 | AF700 or Biotin | BioLegend | HM36 | 100x |
|  | streptavidin-BV650 | BioLegend |  | 400x |
| CD19 or | ef450 | ThermoFisher | eBio1D3 (1D3) | 100x |
| B220 | BV510 | BioLegend | RA3-6B2 | 100x |
| ***Intracellular stain*** |  |  |  |  |
| Glut1 | BSA/Azide free | Abcam | EPR3915 | 200x |
| (Lightning link) | DL405 |  |  |  |
| PKM | BSA/Azide free | Abcam | EPR10138(B) | 1.000x |
| (Lightning link) | PE |  |  |  |
| SDHA | BSA/Azide free | Abcam | EPR9043(B) | 1.000x |
| (Lightning link) | AF647 |  |  |  |
| CPT1A | BSA/Azide free | Abcam | EPR21843-71-2F | 200x |
| (Lightning link) | PE/Cy5 |  |  |  |
| ACC1 | BSA/Azide free | Abcam | EPR23235-147 | 200x |
| (Lightning link) | AF488 |  |  |  |
| Cytc | BSA/Azide free | Abcam | 7H8.2C12 | 1.000x |
| (Lightning link) | PE/Cy7 |  |  |  |
| G6PD | BSA/Azide free | Abcam | EPR20668 | 1.000x |
| (Lightning link) | APC/Cy7 |  |  |  |
| **Surface stain 2** |  |  |  |  |
| CD45 | APC/Fire810 | BioLegend | 30-F11 | 2.000x |
| CD11b | BUV563 | BD Bioscience | M1/70 | 10.000x |
| F4/80 | BUV661 | BD Bioscience | T45-2342 | *200x |
| CD64 | BV711 | BioLegend | X54-5/7.1 | 100x |
| TIM4 | PerCP-ef710 | BioLegend | RMT4-54 | **10.000x |
| MHCII | BUV395 | BD Bioscience | M1/42 | 3.000x |
| PDL1 | BUV737 | BD Bioscience | M1H5 | 100x |
| PDL2 | BV650 | BD Bioscience | TY25 | 100x |
| Ly6G | SB550 | BioLegend | 1A8 | 1.000x |
| Siglec F | BV480 | BD Bioscience | E50-24440 | 400x |
| Ly6C | PerCP-Cy5.5 | BioLegend | HK1.4 | 1.000x |
| CD3 | BV750 | BioLegend | 17A2 | 200x |
| CD98 | BUV615 | BD Bioscience | H202-141 | 400x |
| CD301 | PE/Dazzle594 | BioLegend | LOM-14 | 400x |
| CD38 | BUV805 | BD Bioscience | 90/CD38 (Ab90) | 400x |

\* 6.000x for PEC, \*\* 400x for intestines

## Supplementary Figure Legends

**Figure S1.**
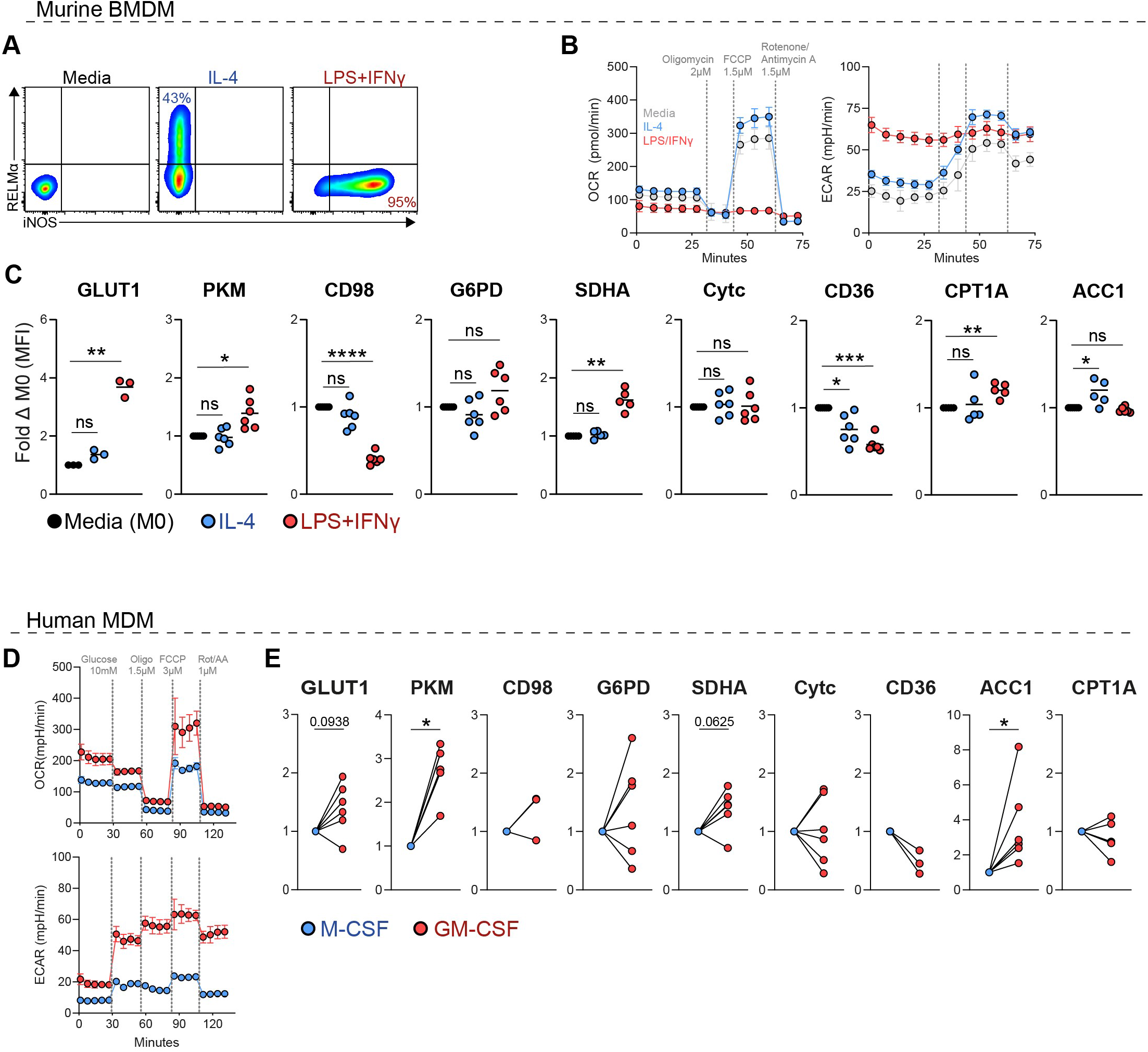
Metabolic flow cytometry identifies distinct phenotypes between in vitro-stimulated macrophages. (**A**) Representative expression of inflammatory M(LPS+IFNγ) and alternatively activated M(IL-4) markers following stimulation of BMDM. (**B**) Seahorse analysis using the mitochondrial stress test, showing the oxygen consumption rate (OCR) and extracellular acidification rate (ECAR) of overnight media, LPS+IFNγ and IL-4 stimulated BMDM. Data shows pooled data of BMDM from 2 mice from one individual run. (**C**) Relative change in geometric MFI of metabolic targets by stimulated BMDM relative to media control. Data points represent BMDM cultures from individual mice, pooled from 2-3 experiments with 1-3 mice per experiment. (**D**) Mitochondrial stress-test of human MDM differentiated with either GM-CSF or M-CSF for 6 days. (**E**) Fold change of metabolic target expression (gMFI) in GM-CSF MDM relative to M-CSF macrophages. Statistics calculated with (C) RM one-way ANOVA or (E) Wilcoxon matched-pairs signed rank test. (*p < 0.05, **p < 0.01, ***p < 0.001, ****p < 0.0001, ns=not significant).

**Figure S2.**
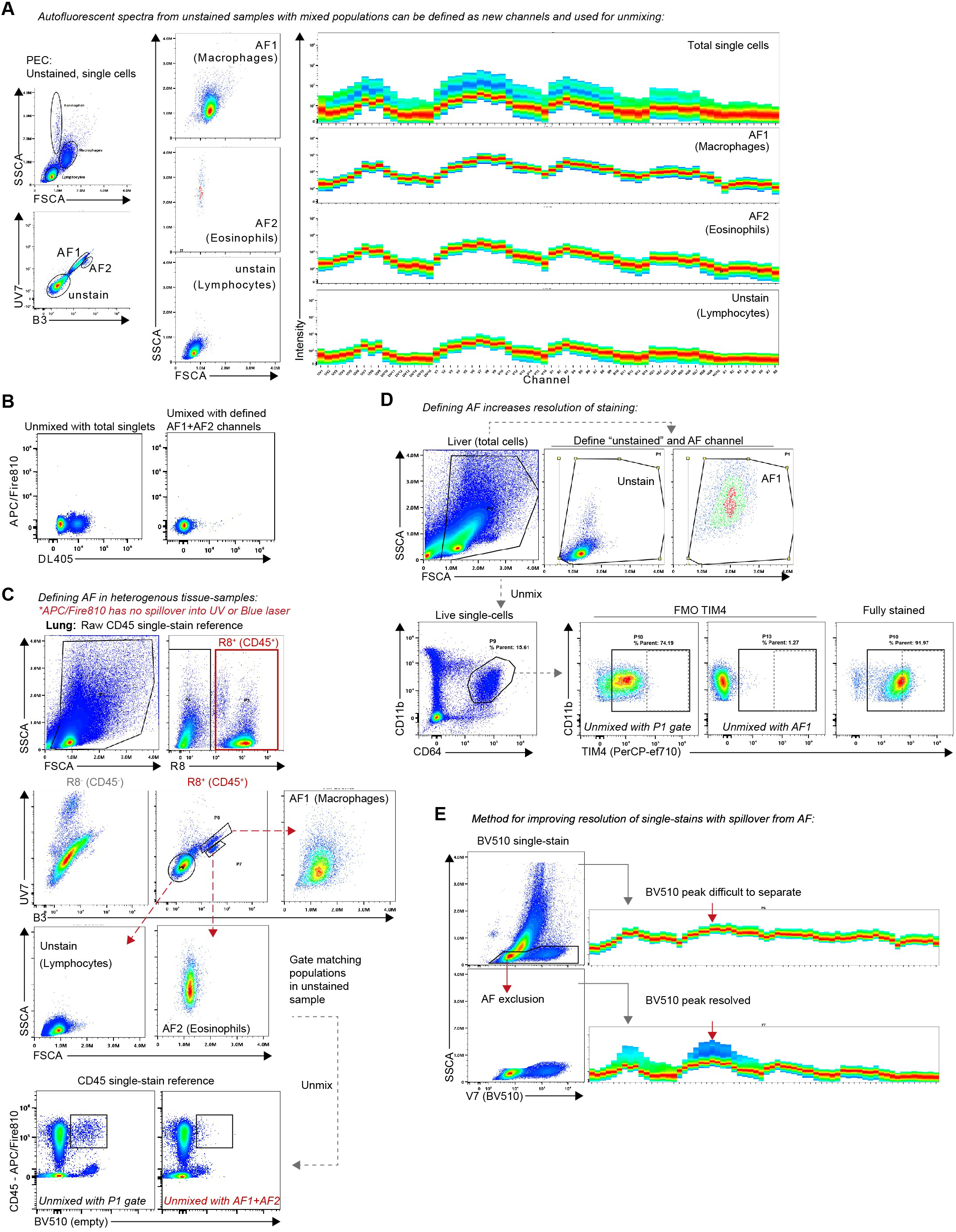
Accounting for autofluorescence in spectral unmixing of heterogenous samples. (**A**) Distinct scatter profiles and spectral signatures of mphs, lymphocytes and eosinophils in the murine peritoneal cavity. (**B**) Representative unmixing of unstained sample for a channel with high similarity to AF (DL405), and a channel with low similarity to AF (APC-Fire810) unmixing with or without AF channels defined in (A). (**C**) Identification of immune cell AF signatures in stromal tissues using raw worksheet, with the lung as an example, and representative unmixing of unstained samples for BV510 (high AF overlap) with and without AF definition. (**D**) Representative unmixing results for TIM4-PerCP-eFluor710 staining on liver mphs with and without defining extra AF channel. (**E**) Example of increasing resolution of single-stained samples by gating out AF before unmixing.

**Figure S3.**
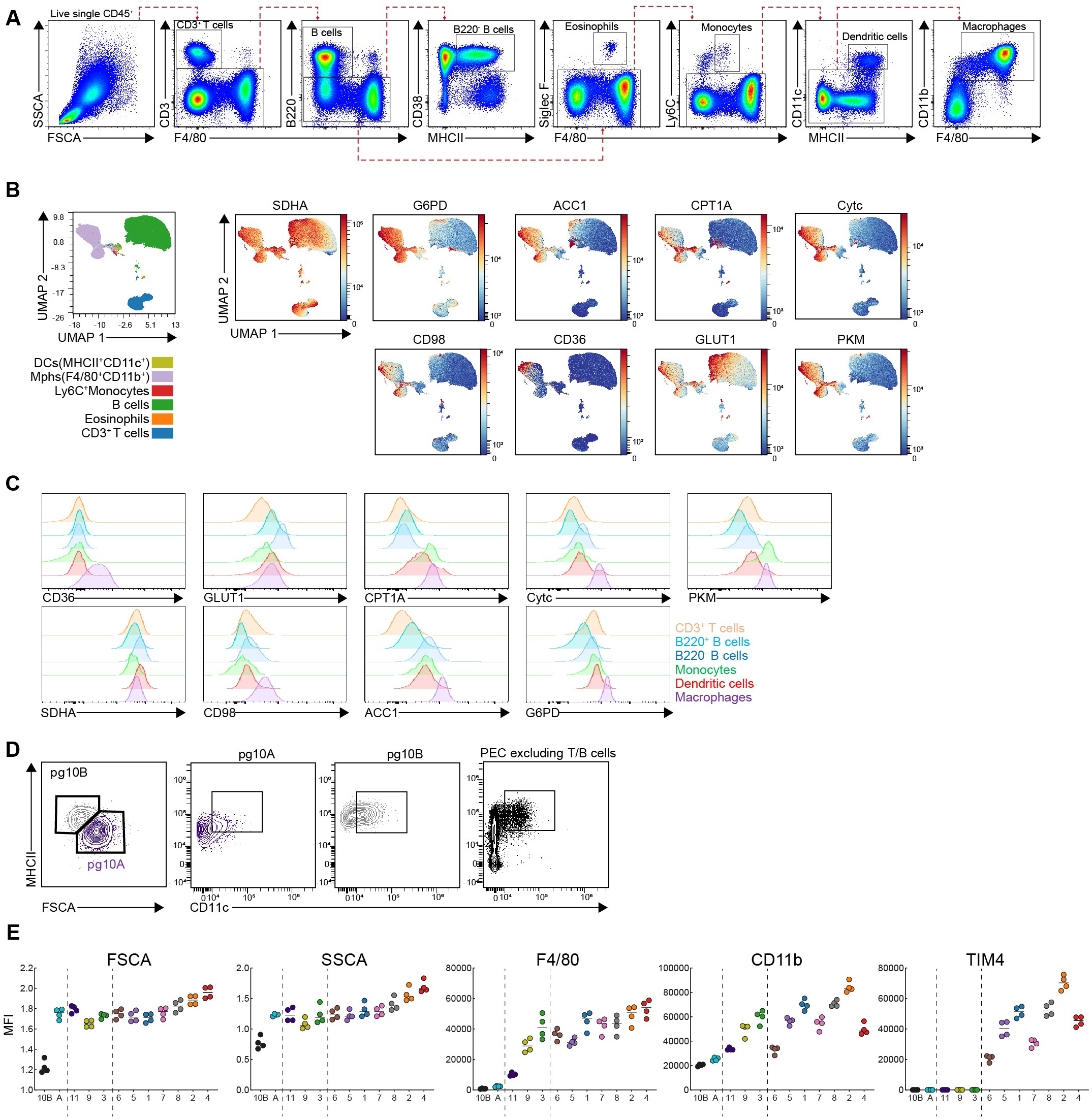
Metabolic heterogeneity of the peritoneal cavity immune compartment. (**A**) Gating strategy for different peritoneal immune cell populations. (**B**) UMAP and phenograph clustering on total immune cells from the PEC, of 4 mice pooled. (**C**) Histograms showing expression of metabolic targets across populations. (**D**) MHCII, FSC-A and CD11c expression in pg10, used to define sub-clusters pg10A and 10B. (**E**) MFI of scatter parameters and lineage markers corresponding to OMIQ analysis in Figure 1.

**Figure S4.**
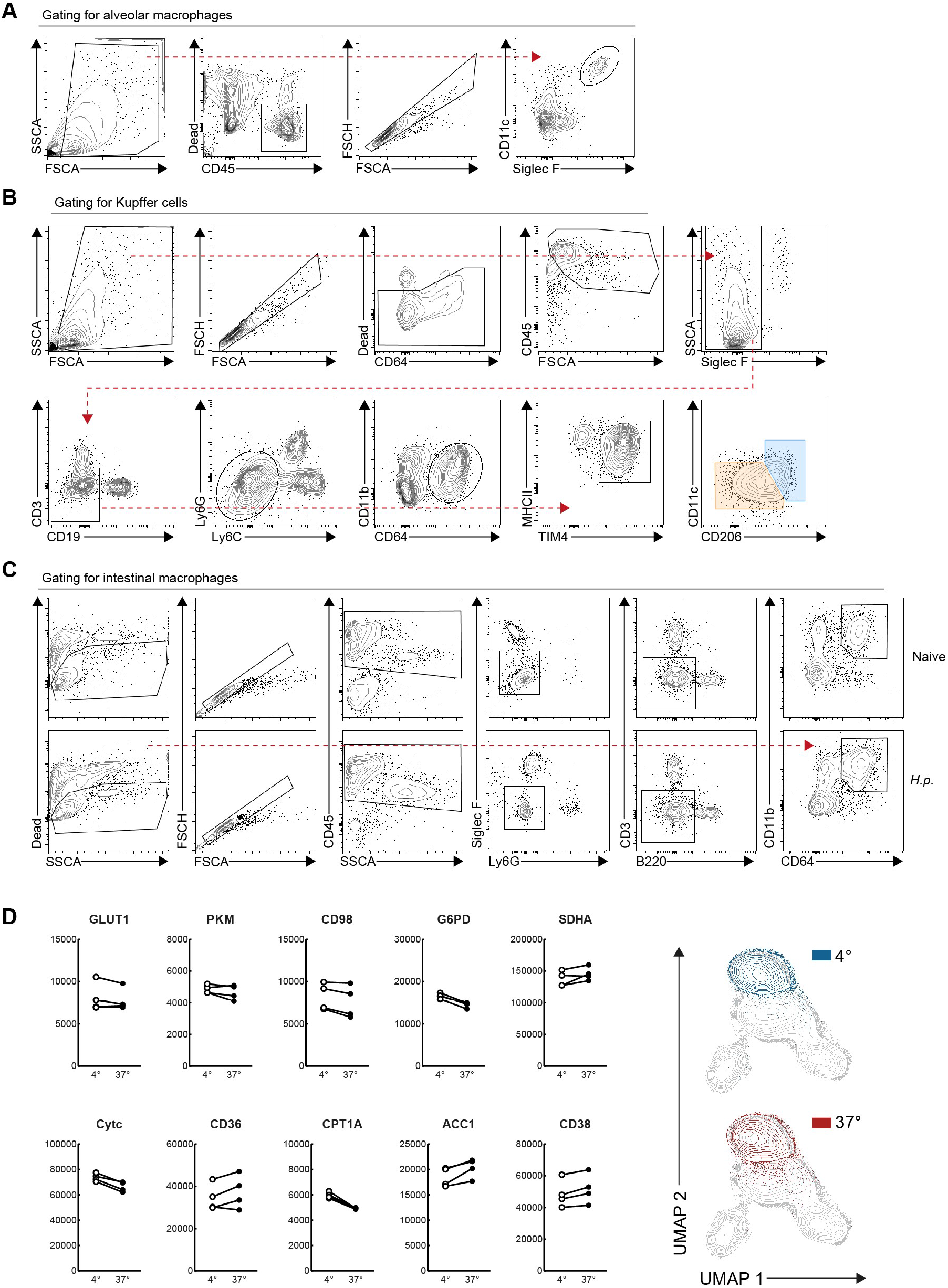
Gating strategies for digested tissues. Representative staining an gating strategies for (**A**) Alveolar mphs, (**B**) Liver Kupffer cells, and (**C**) mphs from the small intestine (duodenum) of naïve and Hp infected mice. (**D**) PEC samples were divided with half stored on ice and the remaining “digested” in RPMI shaking at 37^°^C for 30minin parallel to digesting tissues, then assessed for metabolic expression and overlap following UMAP using metabolic targets.

## Notes

### Competing Interest Statement

The authors have declared no competing interest.

